# A benchmarking model for validation and standardization of traction force microscopy analysis tools

**DOI:** 10.1101/2020.08.14.250506

**Authors:** Gaspard Pardon, Erica Castillo, Beth L. Pruitt

## Abstract

Traction Force Microscopy (TFM) has become a well-established technique to assay the biophysical force produced by cells cultured on soft substrates of controlled stiffness. However, experimental conditions as well as computational implementations can have a large impact on the analysis results accuracy and reproducibility. While this can be alleviated using appropriate controls and a rigorous analytical approach, the comparison of results across studies remains difficult and there is a need for validation and benchmarking tools. To validate the accuracy of and compare various computational TFM analysis algorithms, we developed a virtual *in silico* model of a cell contracting on a soft substrate of controlled stiffness. The model utilizes user-defined parameters for the cell dimensions as well as for the strength and spatial distribution of a contraction dipole to calculate the deformation that would result on a soft substrate due to the cell contraction. The deformation is computed using the forward analytical stress-strain tensor calculation in the Fourier space. The resulting displacement field is used to apply, using image processing, a deformation on a real or simulated image of fluorescent microspheres embedded into a soft hydrogel, which is normally obtain experimentally by TFM imaging. The deformation field and resulting image then serve as input in the PIV and TFM analysis. The model also enables to create movies of a dynamic cell contraction, such as that of a cardiomyocyte, to validate the time accuracy of the TFM analysis after application of image processing algorithm, such as denoising. Our tool therefore addresses the need for validation and standardization of TFM analytical algorithms and its experimental implementations.

## Introduction

Traction force microscopy (TFM) is a well-established tool to measure the biophysical forces exerted by cells on soft substrates. ^1,2^ It relies on the imaging the displacement of fiducial markers, often fluorescent microspheres embedded in a biocompatible hydrogel of controlled stiffness, to compute the traction stress. In TFM, low image quality or inadequate processing parameters can lead to inaccurate computation of traction stress, especially on stiffer substrates and with noisy images, where bead displacement is small and more challenging to measure. It is therefore crucial to quantify the overall accuracy of the computation, as well as the effect(s) of individual processing parameters and of experimental conditions on the results. Further, to render results from various studies comparable, is important to be able to benchmark the analytical algorithm used.

Here, we developed an *in silico* model that addresses this need. Our model was developed in Matlab and enables to generate virtual images or video of the contraction of a cell, using as input a single computer-generated or real microscopy image of randomly distributed fluorescent microspheres as well as user-defined cell geometry and contractile stress. This model is built upon Jorge-Peñas et al work.^3^ The code is available from our Github page.

## 1. Methods

Our model defines a virtual cell and associated force dipole, for which the dimension, orientation, amplitude, area of application, and temporal dynamics can be controlled by design. Using the forward mathematical tensor for the stress-strain relation, the model computes the deformation field that such a force dipole would dynamically induce in a deformable material of the chosen stiffness.^1,4^ The calculated deformation field is then imprinted onto the input image through a warping-function image processing to yield a virtual image or video of a hydrogel under dynamic deformation from a cell (**Figure 1**). To investigate the effects of processing parameters on computational output, we used our model to design videos of a virtual ellipsoidal cardiomyocyte (CM) cell of length 100 μm and width 18 μm, at orientations ranging from −90° to +90°. The virtual CM was designed to produce a contraction dipole total force of maximum amplitude 0.5 μN typical in CMs differentiated from human induced pluripotent stem cells, applying a spatial surface stress distributed as a 2D Gaussian of radius 5 μm at either end of the cell body.

**Figure 1:**
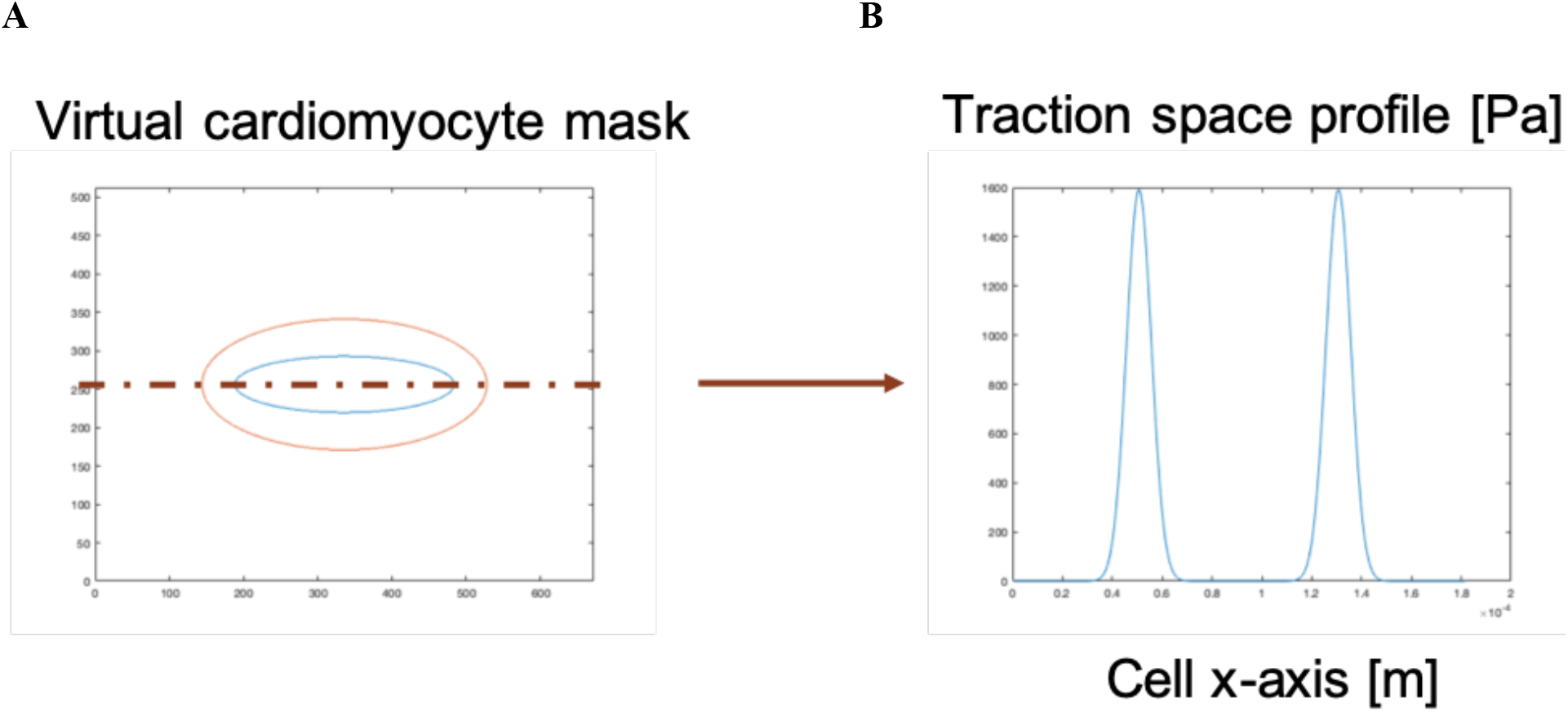
A virtual cell is defined as a ellipse of elongated aspect ratio (A) that produces a contracting force dipole (B).

The force amplitude was applied as a time-varying signal following a Gaussian, triangular, or square signal profile. The exact displacement generated by our model dipole was computed with the exact forward mathematical tensor. The displacement fields were used to apply an image warping transformation to the single static image of the fluorescent microspheres in subsequent steps to generate virtual videos of a contracting cell (**Figure 2**).

**Figure 2:**
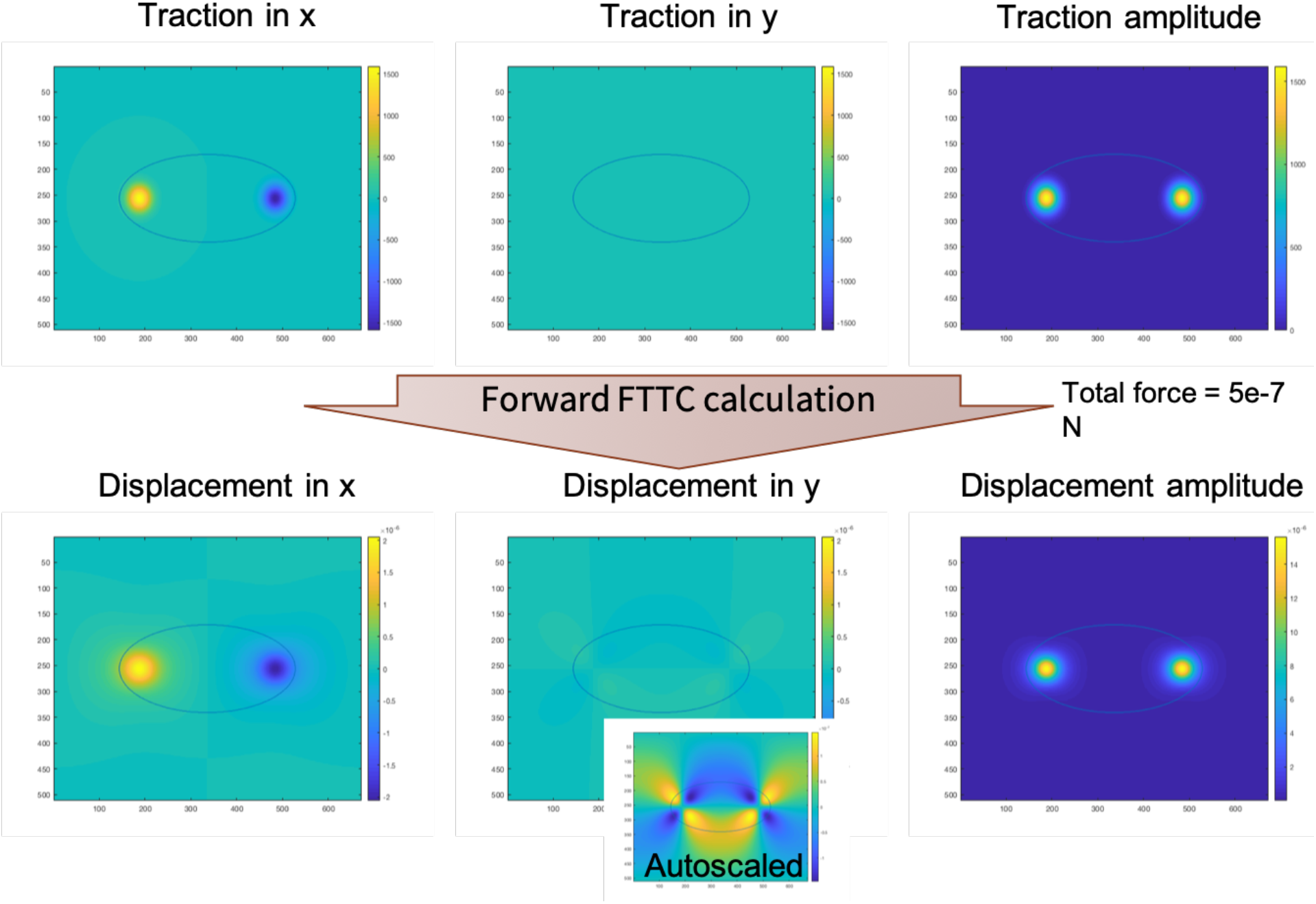
The traction force dipole (Top) is used as input in the model to compute the exact hydrogel deformation displacement (Bottom). Inset in y displacement shows the displacement in the y-direction with tight color scale. This calculated displacement is used to deform the static image of the fluorescent microsphere.

**Figure 3:**
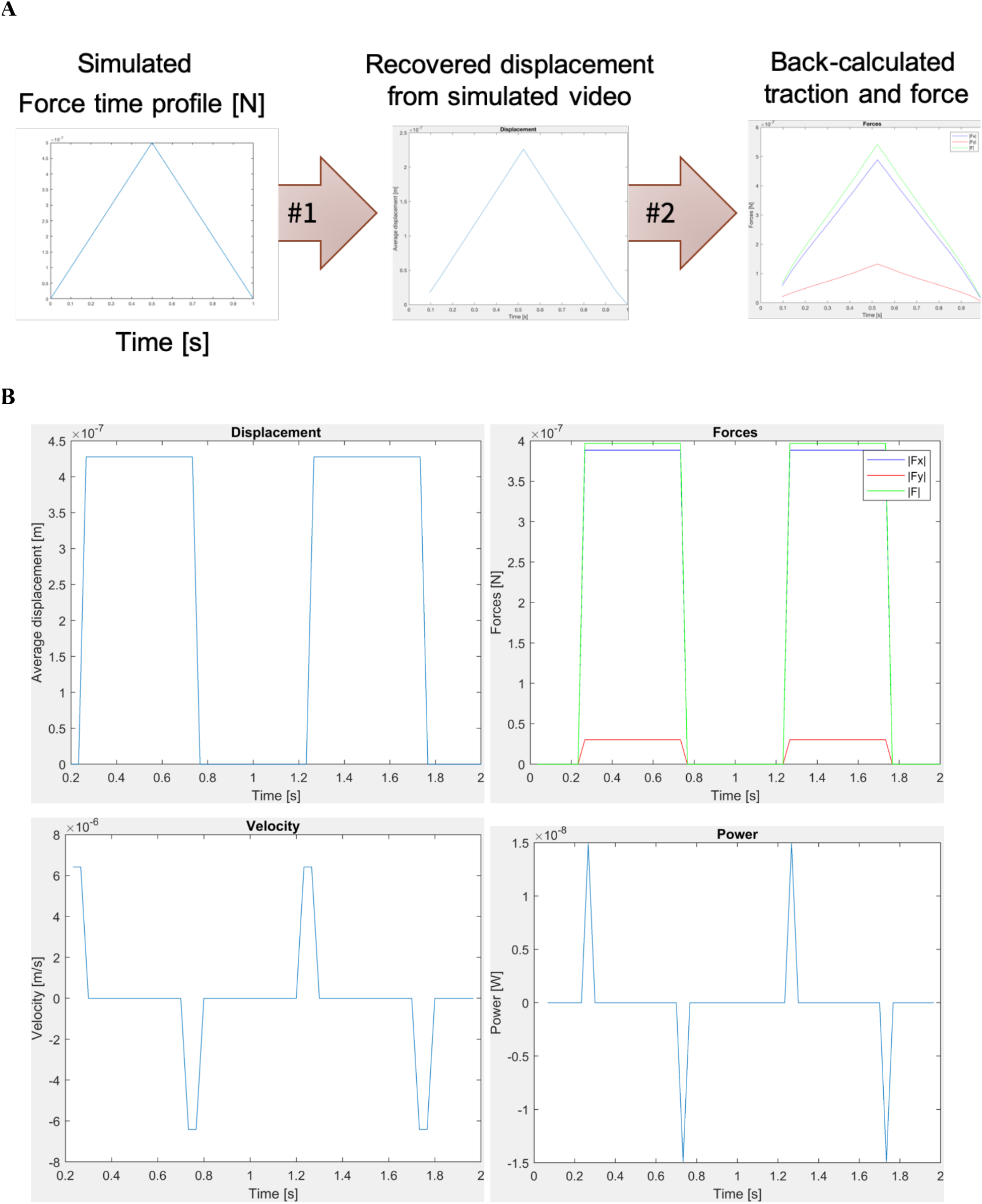
The temporal profile of the force dipole is user defined as a triangle (A), square (B) or following a user-define function (not shown). **A)** The displacement and force are accurately recovered in time. **B)** The square signal is accurately recovered without distortion in time. The force amplitude within 20% of the input. The velocity and power capture the abrupt change as single point peaks.

The displacement field and virtual videos were provided as input to a TFM computation software developed by our lab.^5^ Our computational tool makes use of the NCorr algorithm for the calculation of the fluorescent microsphere displacement by image cross-correlation and of Fourier Transform Traction Cytometry for the reconstruction of the traction stress in the Fourier space, as proposed by Butler *et al.* and Sabass *et al.*. ^1,2^

## 2. Results

The total force recovered was plotted as function of the cell orientation and of the processing parameters for the displacement calculations **Figure 4**.

**Figure 4.**
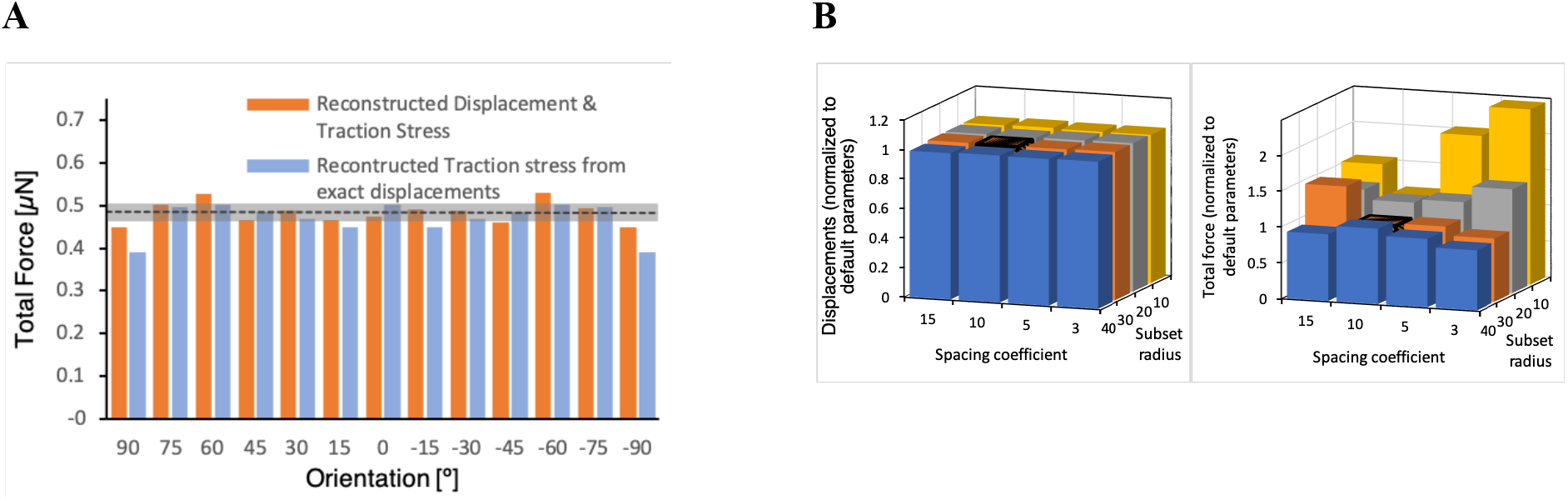
Cell orientation and analytical parameters impact on the accuracy of TFM analysis. **A)** Total force recovery at slightly differs at various cell orientations when computed directly from the simulated displacement or from the virtual video using the TFM algorithm developed in our lab.^**5**^. **B)** PIV processing parameters impact on the accuracy of the calculation of average microsphere displacement and total force.

For our default process parameters, the total force recovered by our TFM analysis algorithm in within an average of 96.8% of the input force when reconstructing traction stress and microsphere displacement from the computer-generated model video, and an average of 93.6% when reconstructing traction stress directly from the exact deformation field computed with the model (Figure 4 **A**). For a horizontally oriented cell, the force recoveries are 94.9% and 100.2%, respectively. A variation of 5.4% results from changing the orientation of the cell in the image frame. There are two explanations for latter variations. First, digitizing the analysis may induce and propagate digital errors due to pixelization. Second, edge effects are known to impact the calculations of microsphere displacement and force; these effects are more pronounced the closer a cell end is to the edge of the image. In such cases, the numerical noise induced by the edge propagates far enough to be included in the surface integration of the traction stress within the cell boundary, impacting the results. We observe that for a cell that is oriented along the diagonal of the frame, i.e. 45°, the recovered force is within less than 1% of the input, since the distance to the image edges is minimal in this configuration.

Varying the analysis parameters for the DIC step (spacing coefficient and subset radius) generates little difference in terms of average displacement within a cell area, but strongly impacts the reconstruction of the traction force (**Figure 4 B**). In our analytical TFM software, DIC is performed using the open-source Ncorr MATLAB package to track hydrogel deformation by following the position of the embedded fluorescent microspheres. In this algorithm, the image is broken down in sub-windows, defined by the subset radius, and interspaced by a specified spacing coefficient, and the position of each window is identified in subsequent frames via 2D cross-correlation. A spacing coefficient smaller than half the subset radius leads to overlapping windows, in effect oversampling the signal, while a larger spacing coefficient leads to undersampling. The size of each window must be defined empirically as a function of experimental conditions (image noise and resolution, microsphere density, and focal depth). Large window sizes underestimate the local displacement by including too large of an area, while small windows sizes yield noisy results. Finally, the smaller the window size and the larger the spacing between windows, the faster the computation, but often at the expense of accuracy. Under our experimental conditions, a spacing coefficient of 10 pixels and a subset radius of 30 pixels yielded the best compromise between results accuracy and computational performance (**Figure 4 B**Error! Reference source not found.).

The code is openly accessible <u>here</u>. ^6^

## 3. Discussion

Our model can also serve as a benchmarking tool for determining or optimizing analysis parameters for specific experimental conditions. Indeed, our model can take as input either a virtual image (for example a dark image of dimensions similar to those of the target video, in which bright dots the size of a fluorescent microsphere are randomly distributed at controlled density) or a real single image of fluorescent microspheres in the hydrogel, allowing benchmarking against a specific imaging setup.

## 4. Conclusion

Overall, our model is a valuable tool to researcher developing TFM algorithms, as it enables validation of the computational accuracy and standardization of results across studies or between laboratories, thereby increasing reproducibility of such assays and increasing the attractiveness of TFM as an approach.

## 5. Author contributions

G.P. and B.L.P. conceived and designed experiments. G.P. developed and designed the computational algorithms and software. G.P. carried out the experiments. G.P., carried out the computation. G.P., E.C with testing of algorithms. G.P., E.C., and B.L.P. analyzed data. All authors contributed scientific insights and to the writing of the manuscript.

## 6. Acknowledgments

We thank all members of the Pruitt laboratory at Stanford University and UC Santa Barbara and to the Blau laboratories at Stanford University for helpful discussions and support.

## 7. Sources of Funding

This research was supported by the Swiss National Science Foundation (SNSF) Early Postdoc Mobility Fellowship (#P2SKP2_164954 to G.P.) and Postdoc Mobility Fellowship (#P400PM_180825 to G.P.), the American Heart Association (AHA Award 18POST34080160 to G.P. and 17CSA33590101 to B.L.P.); the National Institutes of Health (NIH 1R21HL13099301 and RM1GM131981 to B.L.P.)

## 8. Disclosures

All authors declare no conflict of interest.

## Notes

https://zenodo.org/record/3975545#.XzY-Fy17How

### Competing Interest Statement

The authors have declared no competing interest.

